# Laminar Shear Stress Increases the Cytosolic Pool of PINK1 in Endothelial Cells and Enhances Mitophagic Sensitivity Toward Dysfunctional Mitochondria

**DOI:** 10.1101/2025.06.21.660867

**Authors:** Soon-Gook Hong, Junchul Shin, Jason Saredy, Soo Young Choi, Hong Wang, Joon-Young Park

## Abstract

Phosphatase and tensin homolog (PTEN)-induced putative kinase 1 (PINK1) is an essential molecule in mitophagy process in mammalian cells. Mutation or deficiency of PINK1 has been closely related to several disease conditions. The purpose of this study was to determine PINK1 expression levels and subcellular localization under exercise-mimic laminar shear stress (LSS) condition in human aortic endothelial cells (HAECs) or in exercising mice, and its implication on endothelial homeostasis and cardiovascular disease (CVD) prevention. First, LSS significantly elevated both full-length PINK1 (FL-PINK1) mRNA and protein expressions in ECs. Mitochondrial fractionation assays and confocal microscopic analysis showed reduced FL-PINK1 accumulation on mitochondria with an increase in a cytosolic pool of FL-PINK1 under LSS. Mitophagy flux, determined by a mtKeima probe, decreased with intact mitochondrial morphology and membrane potential under LSS, suggesting that elevated cytosolic PINK1 is not utilized for immediate mitophagy inductions. However, increased cytosolic PINK1 seems to elevate mitophagic sensitivity toward dysfunctional mitochondria in pathological conditions. LSS-preconditioned ECs showed lower angiotensin II (AngII)-induced mtDNA lesions and displayed rapid Parkin recruitment and mitophagy induction in response to mitochondrial uncoupler (CCCP) treatment. Exercise-preconditioned mice, a physiological LSS-enhanced model, showed elevated PINK1 expression in ECs of the thoracic aorta compared to sedentary control. In addition, exercise enhanced AngII-induced mitophagy induction in ECs and reduced AngII-induced mtDNA lesion formation in the mouse aorta. Taken together, LSS increases a cytosolic pool of FL-PINK1, which may elevate the mitophagic sensitivity toward dysfunctional mitochondria in ECs.

## INTRODUCTION

Ample evidence indicates that endothelial mitochondria play a variety of roles in maintaining endothelial homeostasis not only supplying adenosine triphosphate (ATP) via oxidative phosphorylation (OXPHOS) as bioenergetic organelles,^1,2^ but also acting as a signaling interface,^2,3^ and biosynthetic hub.^4,5^ Thus, dysfunctional endothelial mitochondria, manifested by increased mitochondrial reactive oxygen species (mtROS) production, mitochondrial DNA (mtDNA) damage, decreased OXPHOS, imbalance in mitochondrial dynamics, have been implicated in defects in the vasculature, which is known to facilitate the progression of cardiovascular diseases.^6,7^ The results emphasize the necessity of a mitochondrial quality control system for vascular health.

Mitophagy is a selective form of autophagy working to eliminate damaged mitochondria as a quality control system to maintain the requisite number of functional mitochondria to meet the energy demands of the cell.^8,9^ Impairment of mitophagy processes is shown in aging and a plethora of pathological conditions, such as neurodegenerative diseases, myopathies, metabolic disorders, inflammation and cancer,^9^ and cardiovascular diseases (CVDs).^10^

Phosphatase and tensin homolog (PTEN)-induced putative kinase 1 (PINK1) is a type I transmembrane protein composed of 581 amino acids including a putative serine/threonine kinase domain and mitochondrial targeting sequence (MTS) at the N-terminus.^11^ Synthesized full-length PINK1 (FL-PINK1) in the cytosol is imported into the inner mitochondrial membrane (IMM) through translocase of the OMM and IMM (TOM and TIM, respectively) under an intact mitochondrial membrane potential. Subsequently, the MTS at the N-terminus of the imported FL-PINK1 is proteolytically cleaved by matrix processing peptidase (MPP) in the matrix and presenilin-associated rhomboid-like protein (PARL) in the IMM, producing products of ∼54 kDa cleaved forms of PINK1 (ΔN-PINK1).^12,13^ After the N-terminal deletion, PINK1 is released into the cytosol in which the N-terminal phenylalanine exposed by the cleavage is recognized by the N-degron type-2 E3 ubiquitin ligases based on the N-end rule, targeting PINK1 to the ubiquitin-proteasome.^14^ However, upon depolarization of the mitochondria, PINK1 imports and cleavages are reduced, and PINK1 starts to accumulate on the outer mitochondrial membrane (OMM) and phosphorylate basal OMM ubiquitin at Ser65 (pS65-Ub), which recruits Parkin from the cytosol to the OMM because Parkin has a high affinity for pS65-Ub.^15^ The recruitment of Parkin on the pS65-Ub complex of the OMM proteins then ubiquitinates mitochondrial proteins such as voltage-dependent anion channel 1 (VDAC1), mitofusin-1 (MFN1), mitofusin-2 (MFN2), and mitochondrial rho GTPase (Miro),^16^ which provides more ubiquitin substrates for PINK1 to phosphorylate recruiting additional Parkin onto OMM proteins showing positive feedback cycles with an amplification of the signals that drives mitophagy.^17^ Likewise, PINK1 continuously monitors mitochondrial health in the first line of mitochondrial quality control system.^17^

Emerging evidence strongly suggests that exercise, as a non-pharmacological treatment, reduces the progression and development of CVDs ^18^ especially by improving endothelial cell function,^19,20^ but the precise mechanisms are not yet completely defined. Laminar shear stress, known to be elevated in the circulation under exercise conditions,^21–23^ is a factor that links exercise and its beneficial effects on ECs. Exercise-induced LSS is known to maintain mitochondrial homeostasis through inducing mitochondrial biogenesis,^24,25^ attenuating mitochondrial ROS (mtROS) generation ^26^ and mitochondria-dependent apoptosis,^27^ maintaining mitochondrial contents,^24^ and enhancing mitochondrial respiration.^28^ However, the effects of laminar shear stress (exercise mimetic) or exercise on PINK1 expression and its subcellular localization in ECs have not yet been understood.

Therefore, the purpose of this study was to determine PINK1 expression and its subcellular localization under an exercise-mimicking laminar shear stress model in human primary endothelial cells and in the aortic endothelium of exercising mice, and its implication on endothelial homeostasis and CVD prevention.

## RESULTS

### FL-PINK1 mRNA and protein expression were significantly enhanced under laminar shear stress in human aortic endothelial cells (HAECs)

First, exercise-mimicking unidirectional laminar shear stress (LSS) was applied to confluent HAECs using an ibidi *in vitro* flow system for 48 hrs. Full-length PINK1 protein (FL-PINK1, 63 kDa) expression was measured by Western blot. Total eNOS (T-eNOS) expression was measured to confirm that LSS was well applied to HAECs. The HAECs were observed to align perpendicular to flow showing an elongated shape compared to HAECs under static condition displaying a cobble stone shape **(Figure 1A)**. FL-PINK1 expression was significantly enhanced under 5 and 15 dyne/cm^2^ LSS conditions compared to static controls **(Figure 1B and 1C)**. Since PINK1 is known as an important mediator of mitophagy, we initially hypothesized that elevated FL-PINK1 in ECs under LSS may be due to elevated mitophagy flux. To test this hypothesis, we utilized Carbonyl cyanide m-chlorophenyl hydrazine (CCCP), which is synthetic mitochondrial uncouplers and lipophilic weak acids acting as protonophores which are a well-known positive control of mitophagy induction.^29,30^ These compounds can freely move across the lipid membranes allowing the protons to cross the membranes depolarizing mitochondrial membrane potential.^29^ Previous literature has shown that CCCP treatment significantly increases PINK1-Parkin-dependent mitophagy showing an accumulation of PINK1 on the mitochondria.^31–35^ Similar with ECs in response to LSS, CCCP treatment (10 µM, 3 hrs) significantly increased FL-PINK1 expression in HAECs in a time-dependent manner **(Figure 1E and 1F)**. However, a quantitative PCR study revealed that the increase of PINK1 protein expression in ECs under LSS was attributable to enhanced PINK1 mRNA expression, but mRNA expression of PINK1 was not enhanced in ECs treated with CCCP **(Figure 1D)**, implying an elevation of mitochondrial accumulation and a resultant reduction of PINK1 degradation in response to CCCP treatment. The elevation of PINK1 mRNA expression in ECs exposed to LSS was also consistently observed in other studies that have similar experimental design, such as GSE83476 and GSE87534 **(Supplemental figure 1)**. The results suggest that elevation of PINK1 protein levels under LSS may be attributable not to mitophagy induction shown in CCCP-treated ECs but to an elevation of PINK1 mRNA expression.

**Figure 1.**
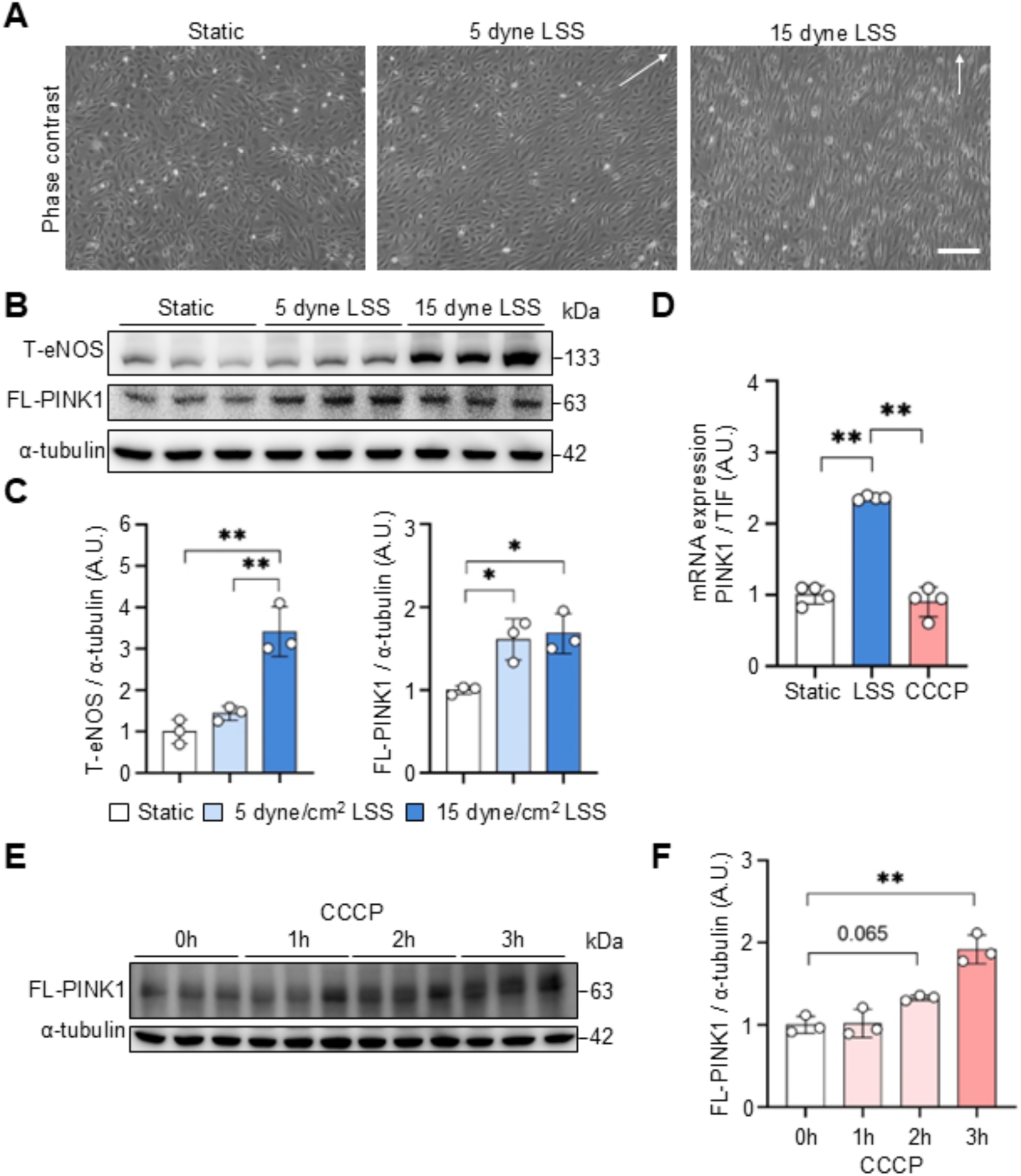
FL-PINK1 mRNA and protein expression were significantly enhanced under laminar shear stress in HAECs. (A) Representative phase contrast images of human aortic endothelial cells (HAECs) under static, 5 dyne/cm^2^ LSS, and 15 dyne/cm^2^ LSS. Arrows represent flow direction. Scale bar = 200 µm. (B) Representative immunoblot images for the protein expression of full-length PINK1 (FL-PINK1) and total endothelial nitric oxide synthase (T-eNOS). alpha-tubulin (α-tubulin) was used as a loading control (n=3/per group). (C) Quantification plots for T-eNOS and FL-PINK1 protein expressions. (D) Quantification plot for PINK1 mRNA expression. Value was normalized by the housekeeping gene eukaryotic translation initiation factor eIF3f (TIF) as an internal control (n=4/per group). (E) Representative immunoblot images for the protein expression of FL-PINK1 and α-tubulin (n=3/per group). (F) Quantification plot for FL-PINK1 protein expression. α-tubulin was used as a loading control. Data is shown as mean ± SD. *p<.05. **p<.01 by one-way ANOVA followed by Tukey’s post hoc test. A.U., Arbitrary Unit.

### LSS increases the cytosolic portion of FL-PINK1 in HAECs

Next, we tested the subcellular localization of PINK1 in HAECs. Mitochondrial and cytosolic fractions were isolated from the HAECs under either static, LSS (15 dyne/cm^2^, 48 hrs), or CCCP treatment (10 µM, 3 hrs), and PINK1 subcellular localization in ECs was analyzed by western blot. As expected, CCCP treatment significantly increases FL-PINK1 accumulation on the mitochondria, which is a known indicator of mitophagy induction. In contrast, LSS decreases PINK1 accumulation on mitochondria but increases the cytosolic portion of FL-PINK1 **(Figure 2A)**. LSS also elevates cleaved PINK1 (ΔN-PINK1) levels, indicating an elevation of PINK1 import and subsequent cleavage due to intact mitochondrial membrane potential (Δψm) under LSS **(Figure 2)**. PINK1 subcellular localization was confirmed by immunostaining and confocal microscopy analysis, which showed that LSS increases cytosolic PINK1 level, but not mitochondrial PINK1 accumulation, showing a reduction of colocalization signals between PINK1 and mitochondria, thereby resulting in a lower Pearson’s correlation coefficient (PCC) compared to static control. In contrast, CCCP treatment enhanced PINK1 accumulation on mitochondria with increased colocalization signals **(Figure 2B and 2C)**. In addition, the percentage of cytosolic PINK1 levels (% of a total PINK1, % Total) was significantly enhanced under LSS compared to static controls **(Figure 2D)**.

**Figure 2.**
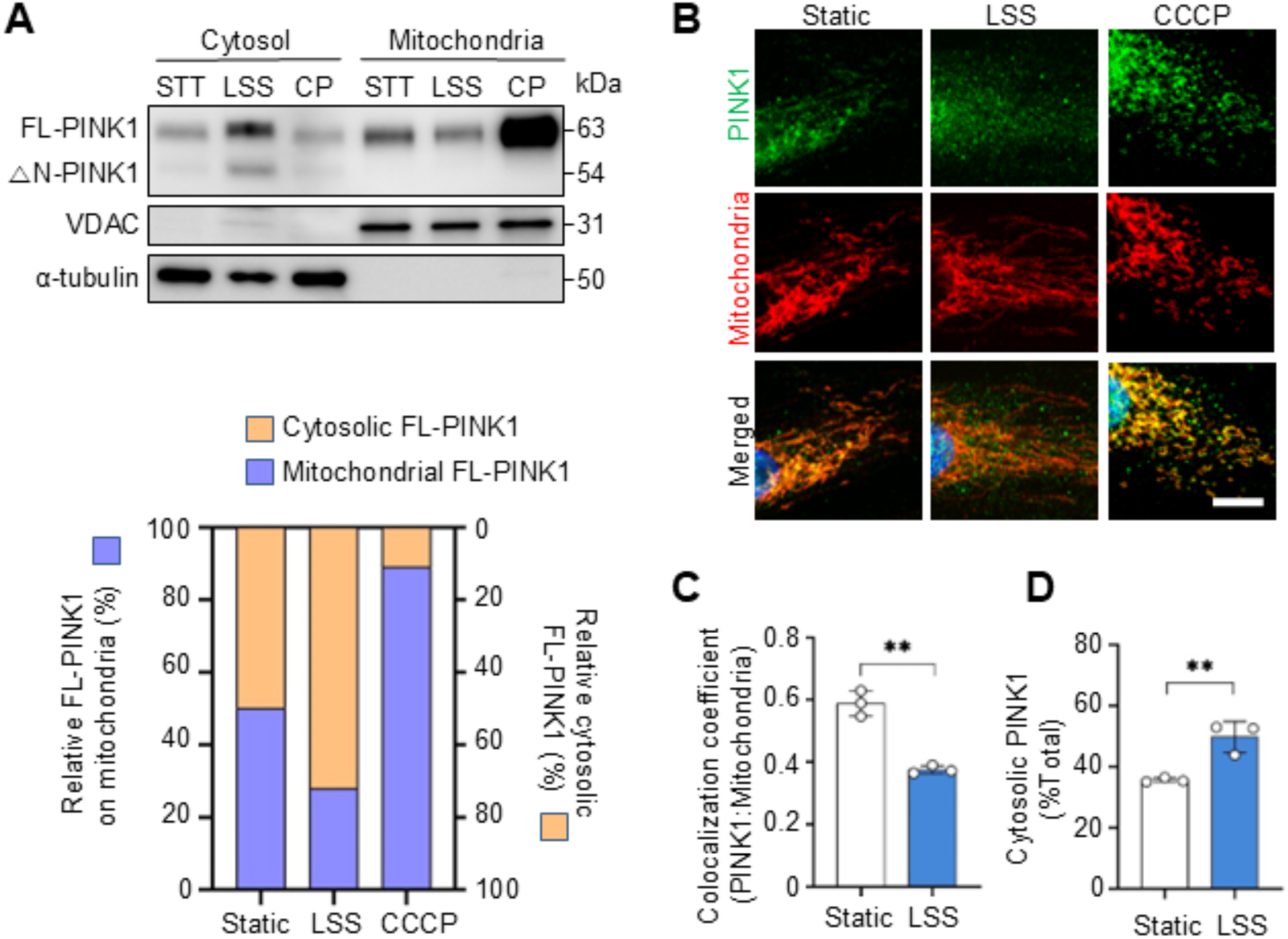
LSS increases the cytosolic portion of FL-PINK1 in HAECs. (A) Representative immunoblot images for FL-PINK1 in the cytosol and mitochondria isolated from ECs under static (STT), 15 dyne/cm^2^ laminar shear stress (LSS), CCCP treatment (CP, 10 µM). Porin (VDAC) and α-tubulin were used as a loading control for mitochondria and cytosol respectively (n=3/per group). (B) Representative confocal images of PINK1 and mitochondria (stained with HSP60) in HAECs under static, LSS, and CCCP treatment (n=3/per group). Scale bar = 20 µm. (C) Quantification plot of colocalization coefficient between PINK1 and mitochondria. Colocalization was quantified from the confocal images by Pearson’s correlation coefficient using ImageJ software. (D) Quantification plot of cytosolic PINK1 (%total). Cytosolic PINK1 was quantified from the confocal images by masking PINK1 signals with mitochondrial signals. Data is shown as mean ± SD. **p<.01 by two-tailed independent Student’s t-test.

### Mitochondria function under LSS is intact with less mitophagy flux

Next, we tested the mitochondrial phenotype under LSS in ECs in that the fate of PINK1 is dependent on mitochondrial function. Initially, mitochondrial morphology was analyzed by confocal microscopy in ECs, which showed that mitochondria displayed elongated and interconnected shapes under LSS (15 dyne/cm^2^, 48 hrs), but fragmented and disconnected mitochondrial shapes were shown in response to CCCP treatment (10 µM, 3 hrs) **(Figure 3A)** resulting in higher mitochondrial fission counts (MFC) **(Figure 3B)**.

**Figure 3.**
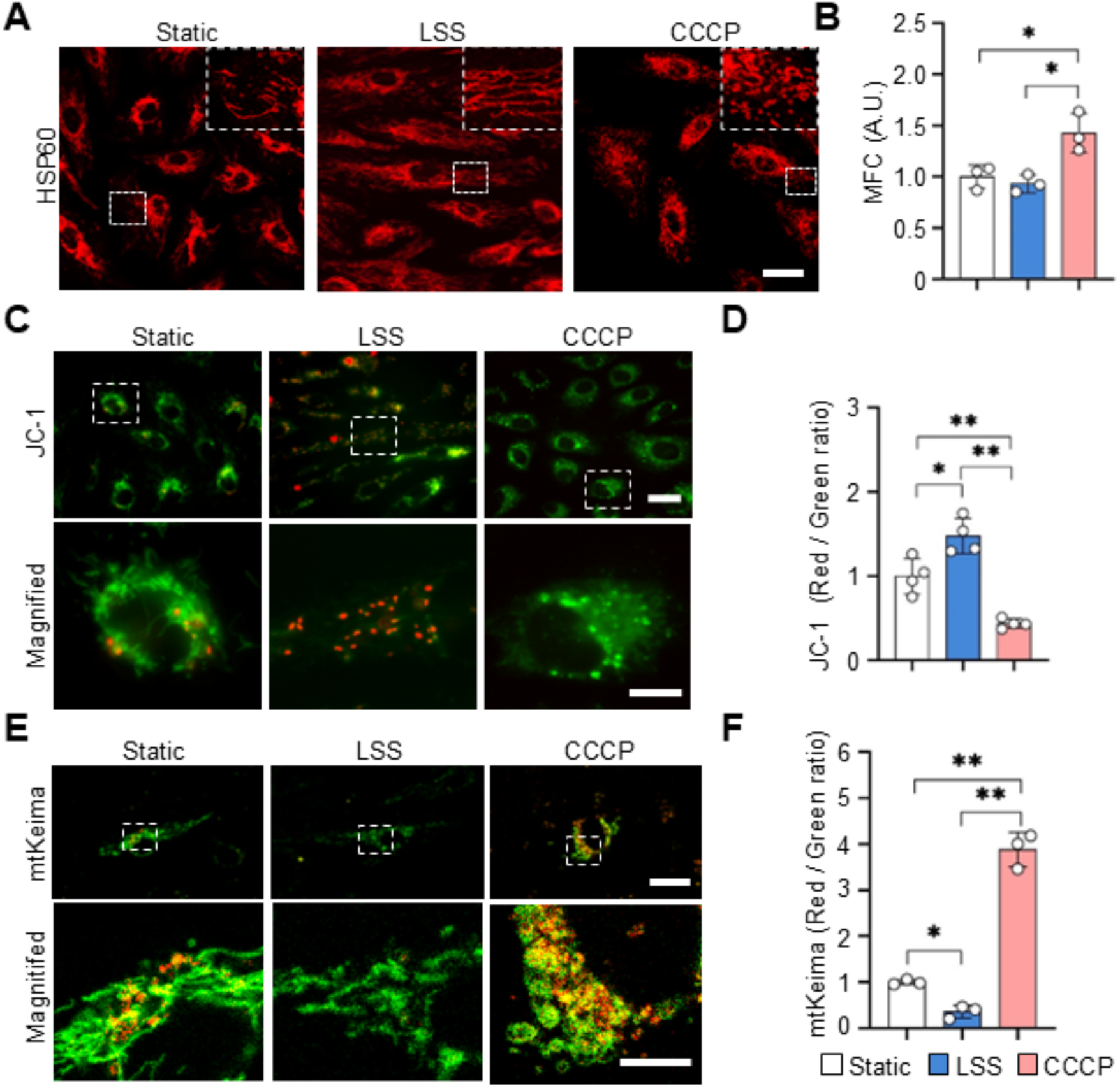
Mitochondria function under LSS is intact with less mitophagy flux. (A) Representative confocal images of mitochondria stained with HSP60 in HAECs under static, LSS, and CCCP treatment. Scale bar = 30 µm. (B) Quantification plot for mitochondrial fission count (MFC = mitochondria number / mitochondrial area) (n=3/per group). (C) Representative fluorescence microscopic images of JC-1 staining in HAECs under static, LSS, and CCCP treatment. Scale bar = 30 µm (upper) and 10 µm (lower). (D) Quantification plot of JC-1 signal by calculating red/green signal ratio (n=4/per group). (E) Representative confocal image of mtKeima in HAECs under static, LSS, and CCCP treatment. Mitophagic flux was monitored in HUVECs transfected with mtKeima plasmid. The red signal represents mitochondria under a mitophagy process, and the green signal represents mitochondria under a normal condition. Scale bar = 30 µm (upper) and 10 µm (lower). (F) Quantification plot of mtKeima signal. The red/green ratio was calculated (n=3/per group). Data is shown as mean ± SD. *p<.05. **p<.01 by one-way ANOVA followed by Tukey’s post hoc test. A.U., Arbitrary Unit.

Based on the morphology analysis, we hypothesized that mitochondrial function will be different under these conditions. To test this, mitochondrial function was measured using the JC-1 probe, which tests mitochondrial membrane potential. LSS (15 dyne/cm^2^, 48 hrs) significantly increases the red/green ratio of JC-1, indicating an increase of mitochondrial membrane potential in HAECs. On the contrary, CCCP treatment (10 µM, 3 hrs) completely reduced the red signal with an increase of green signal showing a reduced red/green ratio in HAECs, indicating reduced mitochondrial membrane potential **(Figure 3C and 3D)**. Indeed, quantification of mitophagy flux using human umbilical vein endothelial cells (HUVECs) transfected with mtKeima plasmid showed decreased red/green ratio under LSS (15 dyne/cm^2^, 48 hrs) compared to static control indicating reduced mitophagy flux **(Figure 3E and 3F)**, which is consistent with reduced mitochondrial PINK1 and an elevation of ΔN-PINK1 under LSS shown in **Figure 2A**. Additionally, western blot analysis also revealed that LSS does not enhance mitophagy induction measured by p62 and Parkin expressions as well as ubiquitin-independent mitophagy measured by BNIP3 expression levels, but LSS significantly increase autophagy process measured by an elevation of LC3-II expression levels **(Supplemental figure 2)**. These results imply that FL-PINK1 protein expression increases due to elevated mRNA expression of PINK1 under LSS, while CCCP increases FL-PINK1 protein expression by an elevation of PINK1 accumulation on mitochondria and a resultant reduction of PINK1 degradation due to reduced membrane potential **(Figure 4)**. These results suggest that PINK1 expression increases under LSS, not for an immediate mitophagy induction, but for other functions.

**Figure 4.**
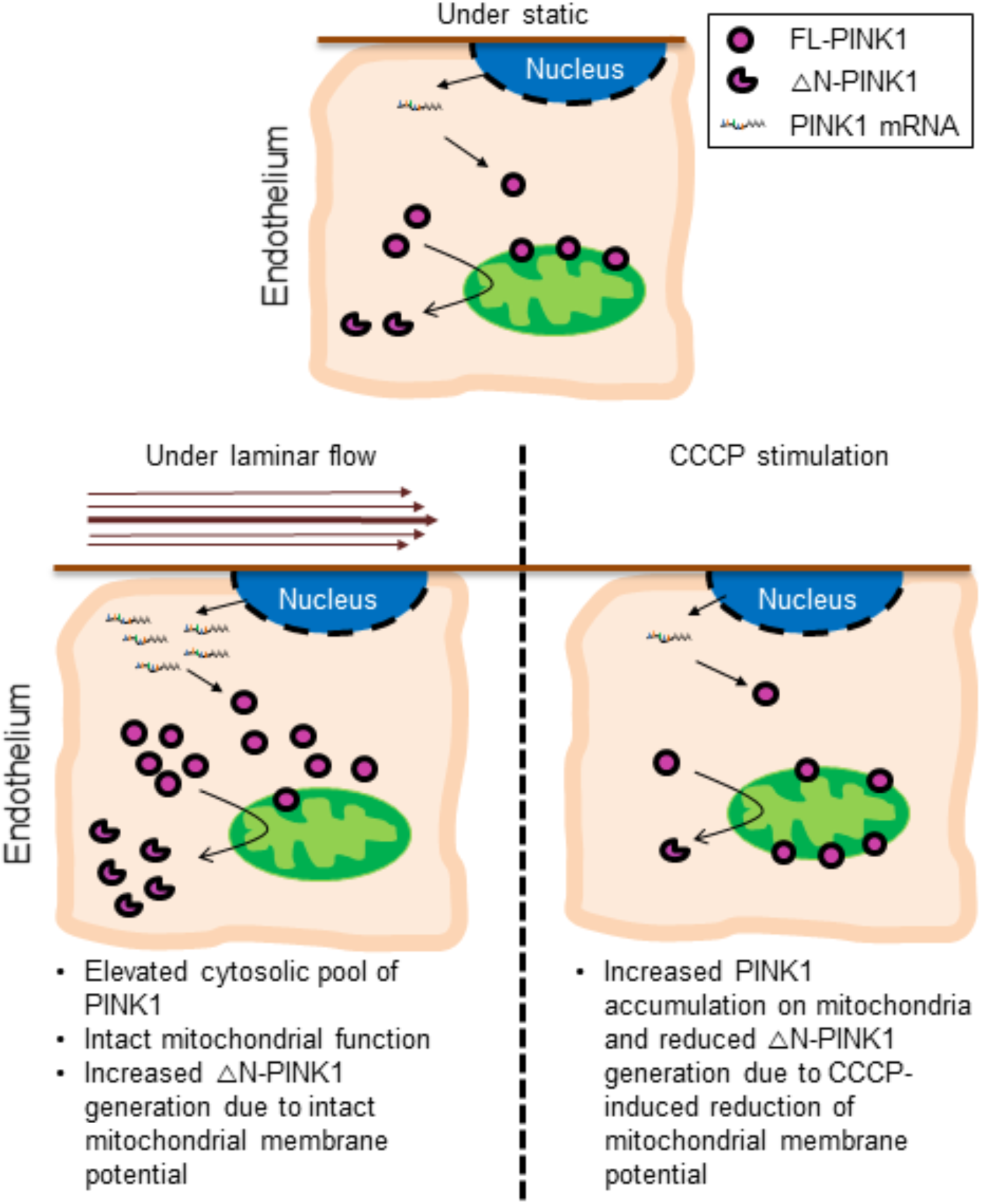
Schematic figure of PINK1 intracellular localization. A cytosolic pool of FL-PINK1 increases under LSS due to elevated mRNA expression of PINK1. In addition, LSS also elevates cleaved form of PINK1 (△N-PINK1) due to intact mitochondrial function under LSS. In contrast, CCCP treatment elevates FL-PINK1 accumulation onto the outer mitochondrial membrane and reduces PINK1 cleavage and degradation due to reduced membrane potential.

### Elevated cytosolic FL-PINK1 may increase mitophagic sensitivity to dysfunctional mitochondria by inducing Parkin recruitment and subsequent mitophagy processes more rapidly to eliminate dysfunctional mitochondria

Based on the observations, we further hypothesized that increased cytosolic pool of the FL-PINK1 under LSS would increase bioavailability of PINK1 to monitor mitochondrial function, which may enhance mitophagic sensitivity to damaged mitochondria under pathological conditions. To test the hypothesis, we preconditioned HAECs under LSS (15 dyne/cm^2^, 48 hrs) for 48h and stimulated them with angiotensin II (100 nM, 6 hrs), which is a well-known peptide hormone that is known to induce mitochondrial DNA damage and eventually their dysfunction. Mitochondrial DNA damage was measured by a mtDNA lesion assay (number of lesions per 10 kb) **(Figure 5A)**. Angiotensin II treatment significantly increases mtDNA lesions in HAECs compared to controls, but LSS-preconditioning significantly reduced the angiotensin II-induced mitochondrial DNA damage in ECs, implying that LSS-preconditioned ECs are more protective against damaged mitochondria accumulation **(Figure 5B)**. Based on the observation that LSS-preconditioned ECs showed lower mtDNA damage under angiotensin II stimulation in HAECs, we hypothesized that the elevation of cytosolic PINK1 may increase mitophagic sensitivity to dysfunctional mitochondria with an enhancement of mitophagy induction, which may allow ECs to maintain a healthy pool of functional mitochondria for survival. To test this hypothesis, we transfected HUVECs with the mCherry-Parkin plasmid, and those cells were subjected to static or LSS conditions (15 dyne/cm^2^) for 48 hrs. The pre-conditioned HUVECs were treated with CCCP (10µM) for 45 min, and Parkin recruitment was observed and quantified. The result showed that Parkin recruitment in response to CCCP was greater in LSS-preconditioned ECs relative to static-ECs **(Figure 5C and D)**. Next, we tested whether LSS-preconditioned ECs induce mitophagy induction rapidly in response to CCCP treatment compared to ECs under static conditions. HUVECs were transfected with mtKeima plasmid, and those cells were subjected to static or LSS conditions (15 dyne/cm^2^) for 48 hrs. The pre-conditioned HUVECs were treated with CCCP (10µM) for 90 min, and mitophagy induction was quantified by calculating the red/green ratio of the mtKeima signals. Indeed, mitophagy induction was significantly elevated in response to CCCP in LSS-preconditioned ECs compared to static-ECs **(Figure 5E and F)**.

**Figure 5.**
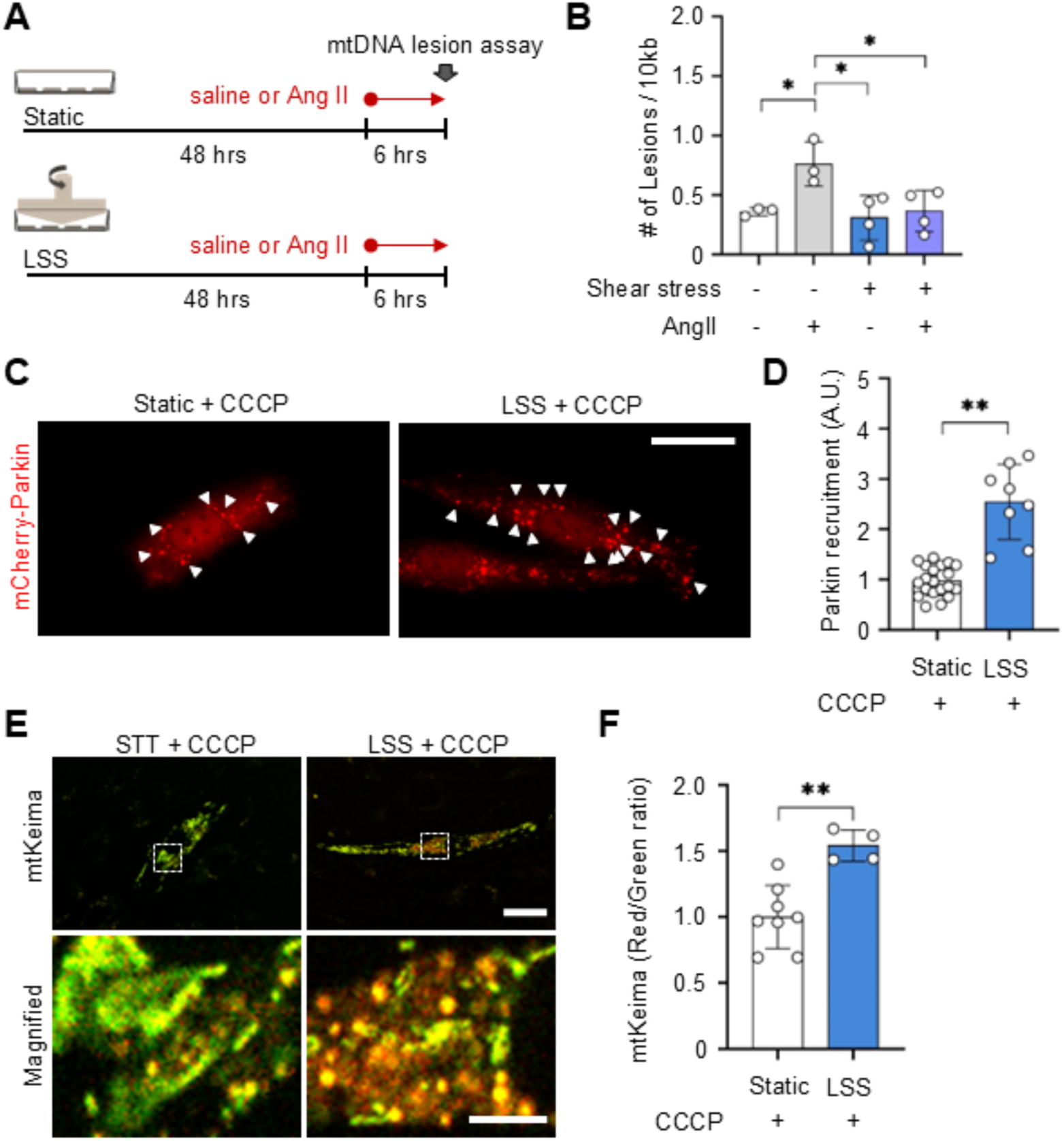
Elevated cytosolic FL-PINK1 may increase mitophagic sensitivity to dysfunctional mitochondria by inducing rapid Parkin recruitment and subsequent mitophagy processes more rapidly. (A) Study design. HAECs were preconditioned under LSS for 48 hrs and stimulated with either saline of angiotensin II (100 nM, 6 hrs) followed by mtDNA lesion assay (# of lesions per 10kb). (B) Quantification plot of mtDNA lesion assay (Number of mtDNA lesion / 10 kb in HAECs) (n=3/per group). (C) Representative micrographs of mCherry-Parkin signal in static– or LSS-preconditioned HUVECs in response to CCCP (10 µM, 45 min). Scale bar = 30 µm. (D) The bar graph is the result of the total area of Parkin recruitment (n=17 for static, n=8 for LSS). (E) Representative fluorescence images of mtKeima signal in static– or LSS-preconditioned HUVECs in response to CCCP (10µM, 90 min). Scale bar = 10 µm (upper) and 2 µm (lower). (F) The bar graph is the result of the red/green ratio of the mtKeima signal. (n=8 for static, n=4 for LSS). Data is shown as mean ± SD. *p<.05. **p<.01 by one-way ANOVA followed by Tukey’s post hoc test (for B) and by two-tailed independent Student’s t-test (for D and F). A.U., Arbitrary Unit.

### PINK1 expression was elevated in the endothelium of thoracic aorta (TA) from EC-PhAM mice subjected to voluntary running wheel exercise in vivo

In order to confirm the observations from the *in vitro* cell studies in mice *in vivo*, we generated endothelial-specific dendra-2 expression mice (EC-PhAM) which shows the endothelial specific mitochondrial fluorescent signals in the mouse blood vessels. The EC-PhAM mice were randomly assigned and subject to either sedentary (n=5) or voluntary running wheel exercise (n=6) for 7 weeks **(Figure 6A)**, and then PINK1 expression in endothelium of the thoracic aorta was measured by *en face* immunostaining **(Figure 6B)**. 7-week voluntary running wheel exercise significantly increases PINK1 expression in the endothelium of EC-PhAM mice, and most of the PINK1 was cytosolic **(Figure 6C and D)**.

**Figure 6.**
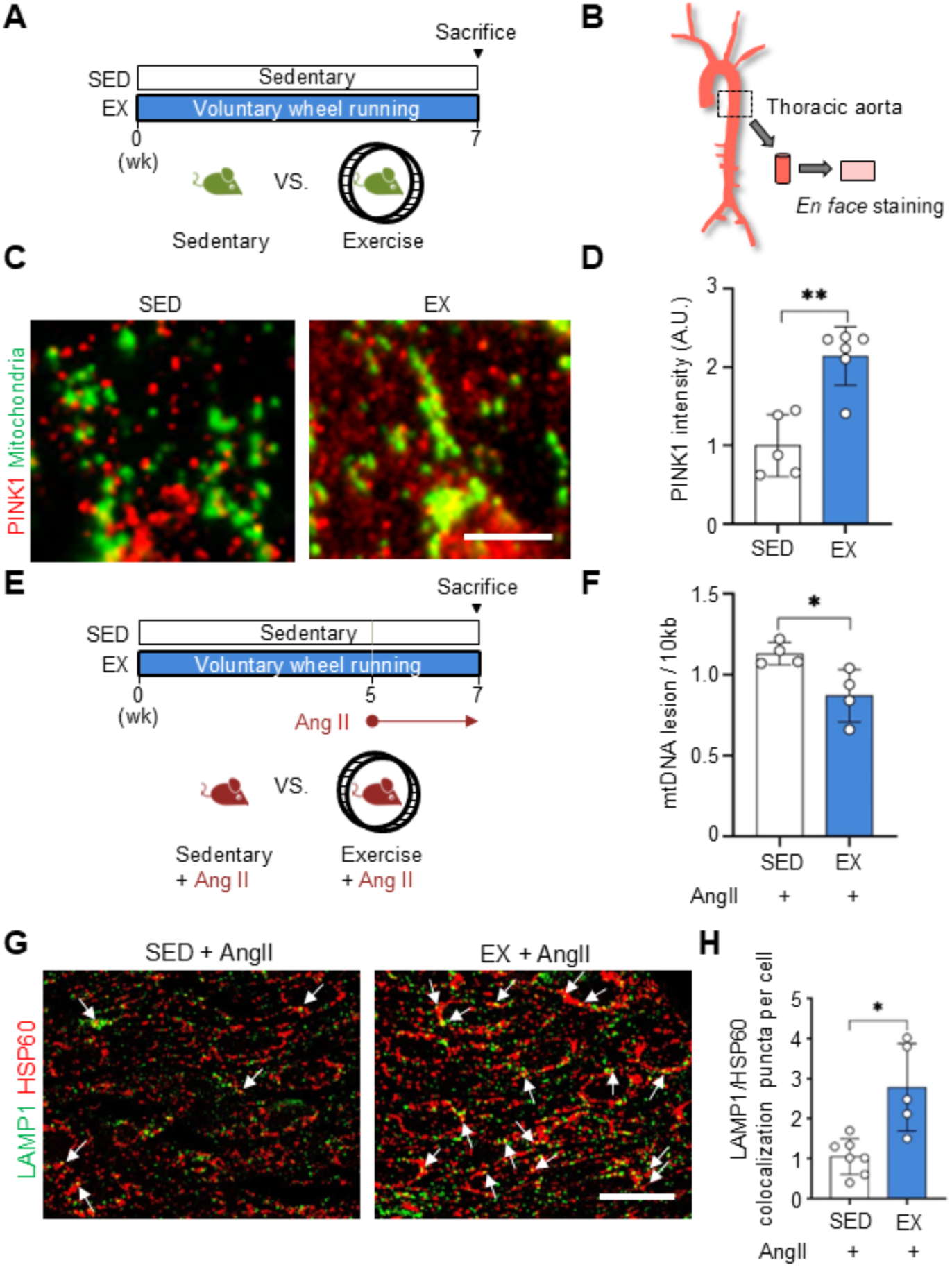
Exercise increases PINK1 expression and elevates mitophagy process in response to AngII stimulation attenuating AngII-mediated mtDNA lesions in the endothelium of mouse aorta. (A) Study design of *in vivo* voluntary wheel running exercise protocol using endothelial-specific Dendra-2 mice (EC-PhAM). (B) Illustration of *en face* staining method. (C) Representative micrographs of PINK1 and mitochondria (mito-Dendra-2) in the endothelium of TA from sedentary (SED, n=5) vs. Exercised (EX, n=6) mice. Scale bar = 30 µm (upper panel) and 5 µm (lower panel). (D) Quantification plot of relative PINK1 intensity in ECs of TA in SED vs. EX. (E) A schematic illustration of experimental design. (F) Quantification plot of mtDNA lesion per 10 kb in mouse abdominal aorta (n=4/group) (G) Representative confocal images of mitophagy induction in endothelium of the thoracic aorta in SED– or EX-preconditioned mice in response to angiotesin II stimulation. LAMP1 (green, lysosome marker), HSP60 (red, mitochondria marker). (H) Bar graph is the number of colocalization puncta between LAMP1 and HSP60 per cell. (n=7 for SED, n=5 for EX). Scale bar = 20 µm. Data is shown as mean ± SD. *p<.05. **p<.01 by two-tailed independent Student’s t-test. A.U., Arbitrary Unit.

### Exercise training elevates mitophagy induction in endothelium of mouse thoracic aorta in response to angiotensin II stimulation and attenuated angiotensin II-induced mtDNA lesion in the abdominal aorta

Next, we tested whether exercise-preconditioned mice show protective effects on mitochondrial health. Mice were randomly assigned and subject to either sedentary (n=7) or voluntary running wheel exercise (n=5) for 7 weeks and stimulated with angiotensin II (1 mg/kg/day) for 2 weeks of total 7-week interventions **(Figure 6E)**. Mitophagy induction and mtDNA lesion were measured in endothelium of the thoracic aorta and abdominal aorta respectively. mtDNA lesion assay showed that exercise-preconditioned mice have less ang-II-induced mtDNA lesions in the abdominal aorta relative to sedentary mice **(Figure 6E and F)**. In addition, exercise-preconditioning elevated mitophagy induction in response to angiotensin II stimulation in endothelium of mouse thoracic aorta measured by colocalization of lysosomal marker LAMP1 and mitochondrial marker HSP60 using *en face* staining **(Figure 6G and H)**.

Based on our data, we concluded that exercise or exercise-mimicking unidirectional laminar shear stress increases a cytosolic pool of FL-PINK1, which may increase mitophagic sensitivity to mitochondrial damage and maintain mitochondrial homeostasis by which exercise may prevent the progression of CVDs on the vasculature **(Figure 7)**.

**Figure 7.**
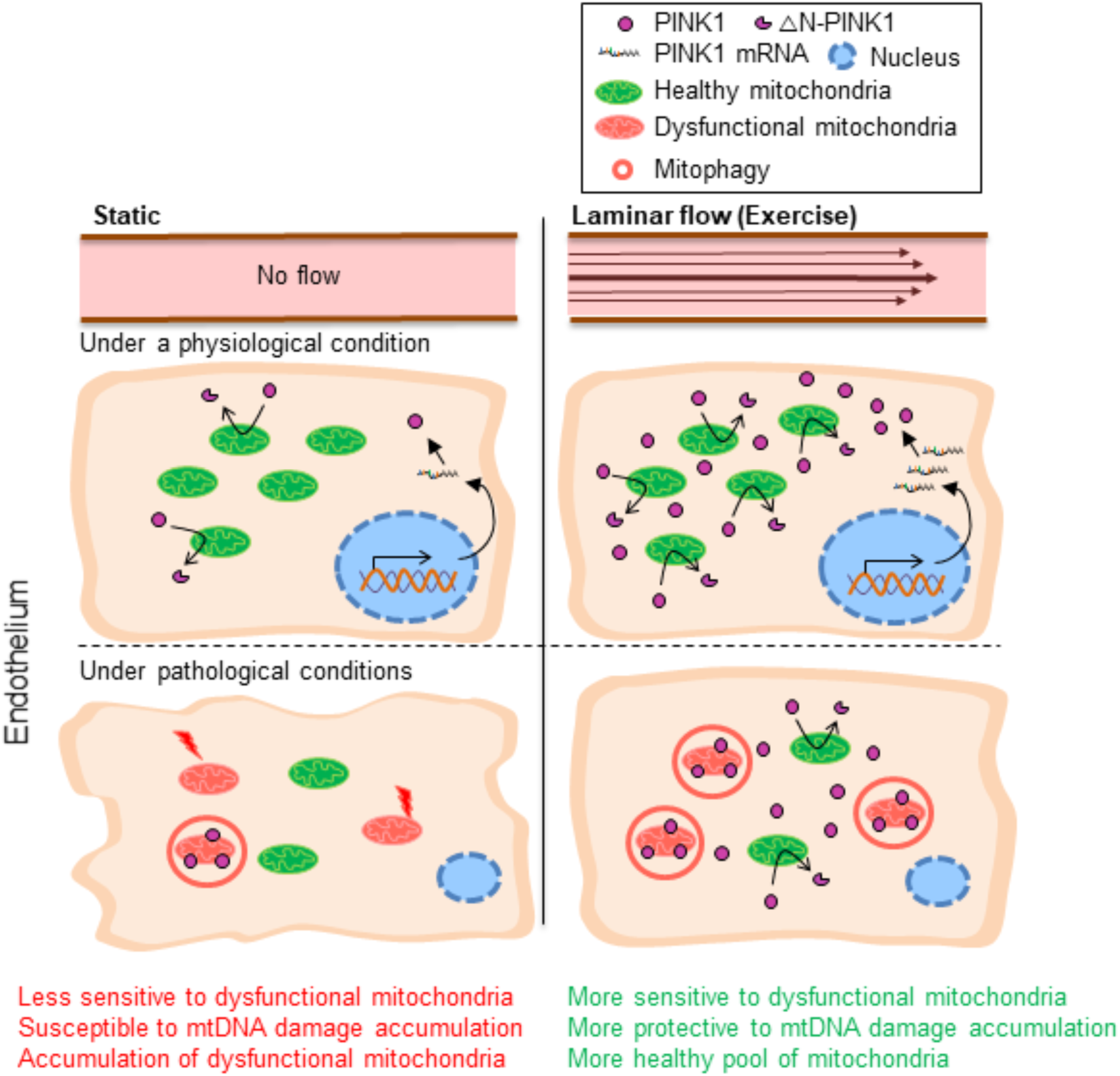
Proposed mechanism of laminar flow-mediated-PINK1 expression and its implication for mitochondrial homeostasis. Laminar shear stress and exercise enhance a cytosolic pool of FL-PINK1, which may increase the mitophagic sensitivity toward dysfunctional mitochondria under stress conditions and thereby render ECs to maintain a healthy pool of mitochondria.

## DISCUSSION

In this study, we demonstrated that LSS increases the cytosolic pool of FL-PINK1, which is attributable to an elevation of PINK1 mRNA expression in ECs. In addition, LSS-preconditioned ECs showed attenuated angiotensin II-induced mtDNA lesions, rapid Parkin recruitment, and mitophagy induction in response to CCCP stimulation, which implies that elevated cytosolic PINK1 expression may enhance mitophagic sensitivity to damaged mitochondria and thereby induce more rapid parkin recruitment and mitophagy induction. Furthermore, exercise training significantly enhanced PINK1 expression in endothelium of mouse aorta *in vivo* in line with *in vitro* observation. The results demonstrate the beneficial effect of exercise-induced LSS on endothelial cell homeostasis by increasing cytosolic PINK1 expression, such that mitophagy can be achieved rapidly before dysfunctional mitochondria accumulate and set up a vicious cycle inducing EC disease-prone phenotype.

Accumulating evidence indicates that PINK1 deficiency is closely associated with CVDs and is known to lead to an accumulation of dysfunctional mitochondria and subsequent cellular defects, whereas overexpression of PINK1 is cytoprotective against various CVD risk factors. Billia et al. (2010) reported that PINK1 protein levels are significantly reduced in end-stage human heart failure, and PINK1^-/-^ mice show development of left ventricular (LV) dysfunction with pathological cardiac hypertrophy displaying greater oxidative stress level and impaired bioenergetic mitochondrial function in mouse hearts.^36^ In addition, Siddall et al. (2013) demonstrated that overexpression of PINK1 in HL-1 cardiac cells was protective toward acute ischemia-reperfusion injuries (IRI), while cardiomyocytes isolated from the PINK1^-/-^ mice showed higher susceptibility to IRI with lower membrane potential and reduced mitochondrial respiration compared to PINK1^+/+^ cells.^37^ Swiader et al. (2016) also reported that silencing PINK1 impaired mitophagy flux and enhanced oxidized LDL-induced apoptosis in vascular smooth muscle cell (VSMC), while overexpression of PINK1 was protective by attenuating VSMC apoptosis.^38^ Similar results have been reported in endothelial cells. Zhang et al. (2014) reported that PINK1-silenced mouse lung endothelial cells (MLEC) show increased susceptibility to hypoxic stress compared to cells from the lung of wild-type mice, while overexpression of PINK1 in MLEC attenuates hypoxia-induced apoptosis in MLEC.^39^ In addition, Wu et al. (2015) showed that mitochondrial damage caused by palmitic acid (PA)-induced metabolic stress was significantly elevated by PINK1 silencing, while overexpression of WT-PINK1 attenuated PA-induced mitochondrial damage measured by mtROS production in HAECs.^40^ Taken together, these findings indicate that PINK1 plays a vital role in maintaining mitochondrial homeostasis and preventing the development of CVDs. In addition, overexpression of PINK1 in different cell types was cytoprotective against a variety of stresses. In line with the PINK1 overexpression studies, our data demonstrated that FL-PINK1 expression was significantly elevated in ECs under LSS compared to static controls **(Figure 1B and 1C)**, and exercise intervention also significantly increased PINK1 expression in the endothelium of the thoracic aorta in mice **(Figure 6C and 6D)**. In addition, LSS-preconditioned ECs and exercise-preconditioned mouse aorta showed an intact mitochondrial pool **(Figure 5B and Figure 6F)** maybe achieved via a PINK1 dependent cellular protection mechanism. Further studies are required to test the casual relationship between LSS-/exercise-induced mitochondrial homeostasis and increased PINK1 expression in ECs.

Previous literature has shown that the autophagy process is significantly elevated under LSS in ECs. Liu et al. (2015) demonstrated that LSS promotes autophagy process via Sirtuin 1 (SIRT1), which subsequently activates FoxO1 by deacetylation increasing its transcriptional activity and thereby enhancing the expression levels of genes related to autophagy processes, such as beclin-1, Atg5, and LC3A.^41^ In addition, Vion et al. (2017) also showed that LSS enhanced autophagy processes in human endothelial cells showing the protective effect on cellular homeostasis. To this end, the deficiency of autophagy in ECs increases endothelial apoptosis, senescence, inflammation, and defects in endothelial alignment leading to EC dysfunction and an atheroprone phenotype.^42^ Furthermore, Li et al. (2015) reported the intact autophagic flux and less mtDNA damage in ECs exposed to LSS both *in vitro* and *in vivo* compared to ECs exposed to atheroprone oscillatory flow.^43^ We also observed an increase of autophagy process measured by LC3-II expression in HAECs under LSS **(Supplemental figure 2)**. Considering enhanced autophagy processes and an increase of PINK1 expression level under LSS, we initially expected an elevation of the mitophagy process under LSS in ECs. However, our data demonstrated that mitophagy flux was reduced under LSS, which was attributed to intact mitochondrial function under LSS in ECs **(Figure 3C)**. In addition, our results from mitochondrial fractionation assay and confocal microscopy analysis supported the observation that elevated FL-PINK1 protein under LSS is not accumulated on mitochondria as an initial step of the PINK1-dependent mitophagy process, but a cytosolic pool of FL-PINK1 was significantly enhanced **(Figure 2)**.

Collectively, these data suggest that increased FL-PINK1 expression under LSS is not enhanced for an immediate mitophagy process due to intact mitochondria under LSS. However, the cytosolic FL-PINK1 may increase bioavailability of PINK1 for more sensitive tests on mitochondrial function and allow ECs to induce resultant rapid mitophagy induction when mitophagy process is required under pathological conditions to eliminate damaged mitochondria before they start to accumulate and set up a vicious cycle leading to an EC dysfunction. Indeed, Narendra et al. (2009) demonstrated that HeLa cells transfected with mCherry-Parkin and PINK1 overexpression vector showed more rapid Parkin translocation to mitochondria in response to CCCP stimulation relative to the cells transfected mCherry-Parkin only, implying the potential role of the elevation of cytosolic FL-PINK1 on mitophagy induction via Parkin recruitment.^33^ Indeed, we showed that LSS-preconditioned HAECs showed rapid Parkin recruitment and mitophagy induction in response to CCCP stimulation **(Figure 5)**, which may be attributable to the elevation of cytosolic FL-PINK1 under LSS in ECs in line with the PINK1 overexpression study.

A large body of evidence indicates that PINK1 can act as a cytoplasmic signaling mediator in the cytoplasm. Indeed, the subcellular location of PINK1 and its role in the cytoplasm has been extensively investigated by previous studies. It has been found that endogenous FL-PINK1 is observed in the cytoplasm ^13,44^ and is known to act as a pro-survival, trophic, and cytoprotective protein.^12,37–39,44–46^ Haque et al. (2007) found that a mutant PINK1 containing a deletion of the putative mitochondrial targeting motif, which is targeted to the cytoplasm, protected neuronal cells from 1-methyl-4-phenylpyridine (MPP^+^)/1-methyl-4-phenyl-1,2,3,6-tetrahydropyridine (MPTP)-induced toxicity.^44^ In addition, Dagda et al. (2014) also reported that transiently transfected neuronal cells with a N111-PINK1 construct that is designed to target PINK1 to the cytosol promoted dendrite outgrowth and restored the decreased dendritic arborization (also known as dendritic branching) by activating PKA (a Ser/Thr kinase) signaling,^47^ which suggests the vital role of cytoplasmic PINK1 in neuronal homeostasis. Furthermore, it has been suggested that PINK1 acts as a cytosolic mediator by utilizing its kinase activity. Murata et al. (2010) revealed that cytoplasmic PINK1 activates mammalian target of rapamycin complex 2 (mTORC2) by phosphorylating a specific component of mTORC2, also known as Rictor, which subsequently phosphorylates Akt (also known as protein kinase B) at Ser-473 showing a protective role in SH-Sy5Y cells from various cytotoxic agents, which suggests that PINK1 can act as a cytoprotective protein not only in mitochondria but also in the cytoplasm through a mTORC2-Akt signaling mechanism.^48^ Therefore, because of the indispensable roles of PINK1, mutation or deficiency of PINK1 leads to the accumulation of mitochondrial defects which eventually results in cellular dysfunction.^45,49^ On the contrary, it has been known that overexpression of PINK1 is cytoprotective in endothelial cells (ECs) against various CVD risk factors.^39,40^ Considering these data showing a cytoprotective role of cytosolic PINK1, increased cytosolic pool of PINK1 under LSS may account for the beneficial role of exercise-induced LSS on endothelial homeostasis. Therefore, future studies are warranted to investigate whether the elevation of a cytosolic pool of PINK1 under LSS modulates cytosolic signaling pathways.

## CONCLUSION

The present study demonstrated that LSS increases a cytosolic pool of PINK1 in ECs, which seems to render ECs more sensitive to dysfunctional mitochondria and thereby induces more rapid mitophagy induction and thereby maintaining a healthy pool of mitochondria in ECs. Our data suggest that exercise may support mitochondrial homeostasis in vascular endothelial cells by enhancing PINK1-dependent cell protection mechanisms.

## METHODS

### Mitochondrial fractionation assay

Mitochondrial fractions were isolated using a mitochondria isolation kit (Thermo Fisher, #89874) following the manufacturer’s protocol. Amicon Ultra-0.5 Centrifugal Filter Unit (Millipore, UFC501096) was utilized to get a concentrated cytosolic protein. Each isolation fraction was lysed with RIPA buffer containing 10 mM Tris-HCl, 5 mM EDTA, 150 mM NaCl, 1% Triton X-100, 0.1% SDS, 1% Deoxycholate, pH 7.5 and centrifuged at 16,000g for 15 min at 4℃. Then, supernatants were collected and subjected to BCA protein assay and subsequent immunoblotting.

### Immunoblotting

Immunoblotting was conducted as previously described.^24^ Briefly, cells were lysed in RIPA buffer, and then collected RIPA samples were centrifuged at 16,000g for 15 min at 4℃. Supernatants were collected and subjected to BCA protein assay (Pierce™ BCA Protein Assay Kit, #23225) to quantify protein concentrations. The protein samples were subjected to sodium dodecyl sulphate–polyacrylamide gel electrophoresis (SDS-PAGE) and transferred to a polyvinylidene difluoride (PVDF) membrane. Subsequently, the membrane was incubated with 5% nonfat dry milk in Tris-buffered saline with Tween 20 (TBST) for 20 min at room temperature (RT) and then incubated overnight with respective diluted primary antibodies. The membrane was washed three times in TBST and incubated with respective HRP-conjugated secondary antibodies for 1 hr. Then, the membrane was washed with TBST and subjected to the standard enhanced chemiluminescence method for visualization.

### Immunostaining

Cells were fixed with 4% PFA for 15 min at RT and washed with PBS three times. After being blocked for 1 hr at RT with staining buffer (10% normal goat serum in PBS containing 0.3% Triton X-100), the primary antibody in the staining buffer was incubated overnight at 4℃. After washing with PBS three times, the secondary antibody in the staining buffer was incubated for 2 hrs at RT. After washing with PBS three times, cells were mounted on DAPI-Fluoromount-G (SouthernBiotech, 0100-20), and images were acquired under a fluorescence microscope (Axioimager, Zeiss) and a confocal laser scanning microscope (SP8, Leica).

### Mitochondrial membrane potential measurement

To measure mitochondrial membrane potential, JC-1 dye (Molecular probes, T3168) was used. Briefly, 10 µg/ml of JC-1 dye in complete media was incubated for 10 min at 37℃ and then washed three times with pre-warmed PBS. Then pre-warmed complete media was added for imaging. Images were acquired under a fluorescence microscope (Axio Observer.Z1, Zeiss)

### Mitochondrial DNA lesion assay

Long Q-PCR was conducted using GeneAmp XL PCR Kit (Applied Biosystems, N808-0192) as described by Janine Santos et al (2006).^50^ Two sets of primer pairs for long Q-PCR are provided below:

> 248 bp short mitochondrial fragment amplicon qPCR primers
>
> Forward: 5’ – CCCCACAAACCCCATTACTAAACCCA – 3’
>
> Reverse: 5’ – TTTCATCATGCGGAGATGTTGGATGG – 3’

> 8.9 kb long mitochondrial fragment amplicon qPCR primers
>
> Forward: 5’ – TCTAAGCCTCCTTATTCGAGCCGA – 3’
>
> Reverse: 5’ – TTTCATCATGCGGAGATGTTGGATGG – 3’

PCR products were quantified by a fluorimetric method using PicoGreen double-strand DNA (dsDNA) quantitation reagent (Invitrogen, P7581), which is known to exhibit over a 1000-fold increase in fluorescence signal when binding to double-stranded DNA (dsDNA). The fluorescence signal was detected using a fluorescence reader with 485 nm excitation and 535 nm emission. Mitochondrial DNA (mtDNA) damage is displayed as lesions per kilobase (kb) mathematically by assuming a Poisson distribution of lesions. Assuming the Poisson equation is defined as f(x) = e^-λ^ λ^x^/x! (zero class, x = 0, molecules exhibiting no damage) and a random distribution of lesions, amplification is directly proportional to the fraction of undamaged DNA templates. Therefore, the average lesion frequency per strand = –lnA_D_/A_O_, where A_D_ represents the amount of amplification of the damaged template and A_O_ is the amplification product from undamaged DNA.^51^

### Animals

All mouse experiments were approved by the Temple University Animal Care and Use Committee (ACUC#4798) and conformed to the NIH guidelines for the care and use of laboratory animals. PhAM floxed mice (B6;129S-Gt (ROSA)26Sor^tm1(CAG-COX8A/Dendra2)Dcc^/J, #018385) and VE-Cadherin-Cre mice (B6;129-Tg(Cdh5-cre)1Spe/J, #017968) were purchased from the Jackson laboratory. To visualize the endothelial mitochondria structure, we bred the PhAM floxed mice with VE-cadherin-Cre mice to generate endothelial-specific PhAM mice (EC-PhAM), which allowed visualization of mitochondria specifically in the endothelium of blood vessels. All mice were fed a chow diet and water ad libitum under a 12-hour light/dark cycle. Angiotensin II (AngII, 1 mg/kg/day) was infused to mice using micro-osmotic pumps (Alzet, #1002) for 2 weeks following 5 weeks of sedentary or voluntary wheel running period.

### en face immunostaining

Mice were perfused with ice cold-PBS with an incision of the right atrium to release the blood followed by a perfusion with ice-cold 2% paraformaldehyde (PFA). The thoracic aorta was isolated and post-fixed with 0.4% PFA overnight at 4℃. The vessel was then washed three times with PBS and incubated with 0.1M Glycine in 2% BSA/PBS for 30 min at RT. Then, the vessel was permeabilized by incubating with 0.3% Triton-X in 2% BSA/PBS for 30 min at RT followed by incubation with primary antibodies in 2% BSA/PBS overnight at 4℃ with gentle agitation. After rinsing in 2% BSA/PBS three times, the vessel was incubated with corresponding secondary antibodies for 2 hrs at RT followed by washing with PBS three times. The vessel was then placed on a slide glass, cut longitudinally, and mounted in DAPI Fluoromount-G (SouthernBiotech, #0100-20). A fluorescence microscope (Axioimager, Zeiss) with a 63X oil objective lens was used for imaging.

### Voluntary wheel running exercise

After the three-day acclimation period, EC-PhAM mice described above (total n=11) were randomly assigned to either the sedentary (SED) (n= 5) or voluntary wheel running exercise (EX) (n= 6) group. EX group animals were individually housed in rat-sized cages with a metal wheel with a diameter of 11.5 cm (Prevue) fitted with a digital magnetic counter for running distance measurement, while SED group animals were singly housed in the same sized cage without the running wheel for 7 weeks. Voluntary wheel running exercise began at 8-week-old and continued for 7 weeks.

### Cell culture and shear stress application

Human aortic endothelial cells (HAECs) (Lonza) and human umbilical vein endothelial cells (HUVECs) (Lonza) were cultured in EGM-2 (EBM^TM^ Basal Medium, Lonza, CC-3121 / EGM^TM^ Endothelial Cell Growth Medium BulletKit^TM^, Lonza, CC-3124) under standard culture condition (37°C under a humidified atmosphere containing 5% CO_2_). All experiments with HAECs were conducted between the 5–8 passages and HUVECs between 4-7 passages. The ibidi pump system (ibidi, Germany) was utilized for applying unidirectional laminar flow (LSS, 15 dyne/cm^2^) to HAECs or HUVECs seeded onto each µ-slide following the manufactural protocol. The perfusion sets and fluidic units were kept and operated in a 37℃ and 5% CO_2_ incubator. When ECs formed a confluent cell layer, LSS was applied to the ibidi μ-slides for 48 hours. For Carbonyl cyanide m-chlorophenyl hydrazine (CCCP, Sigma-Aldrich, #555-60-2) treatment, 10 µM CCCP was treated to ECs once ECs reached 100% confluency.

### Plasmid DNA transfection

mCherry-Parkin was a gift from Richard Youle (Addgene plasmid # 23956; http://n2t.net/addgene:23956; RRID: Addgene_23956).^52^ mKeima-Red-Mito-7 was a gift from Michael Davidson (Addgene plasmid # 56018; http://n2t.net/addgene:56018; RRID: Addgene_56018). The plasmid DNA was purified using the QIAGEN Plasmid Mini Kit (#12123). Transfections of mCherry-Parkin or mKeima-Red-Mito-7 plasmid were performed using Effectene transfection reagent (QIAGEN, #301425) according to the manufacturer’s recommendation.

### Mitochondrial morphology quantification

Mitochondrial morphometric analysis was performed based on the methods described previously.^53,54^ Briefly, obtained images were processed using Image J (NIH) to subtract backgrounds and subjected to kernel convolution (matrix h described below) to emphasize the edges of each mitochondrial particle. Then, images were subjected to binary conversion.

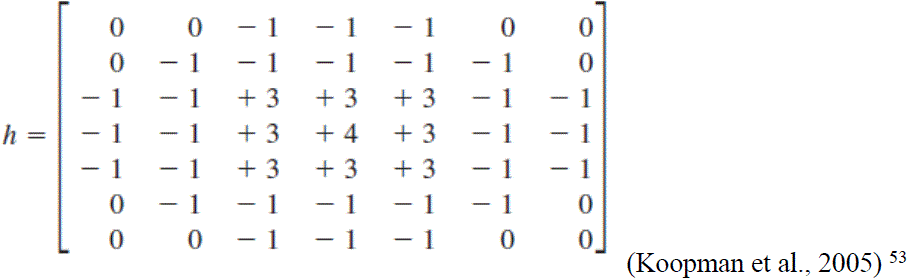

After binary conversion, mitochondrial segments were identified, and continuous mitochondrial structures were counted with the ‘analyze particles’ function of ImageJ, and the number was normalized to the total mitochondrial area to obtain the mitochondrial fragmentation counts (MFC) for each imaged cell (MFC = mitochondria number / total mitochondrial area).

### real-time PCR

Cells were lysed using RNA lysis binding buffer (1M Tris-HCL ph7.5, 5M LiCi, 0.5M EDTA pH8.0, 10% LiDS, 0.1M DTT). mRNAs were isolated with the use of Dynabeads direct kit (Invitrogen, #61011), and cDNA synthesis were performed on poly-dT magnetic beads by reverse transcription using superscript II (Invitrogen, #18064014). mRNA expression levels were quantified by real-time PCR using SYBR green fluorescence. Cycle threshold (Ct) values were normalized to the housekeeping genes. The primer sequences used are described below:

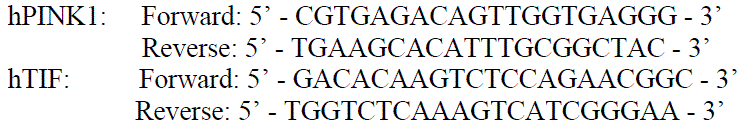

### Antibodies

Primary antibodies used in western blot analysis and immunostaining were as follows: anti-PINK1 (Cell Signaling, #6946) and (Novus, BC100-494SS), anti-VDAC (Porin) (Abcam, ab15895), anti-HSP60 (H-1) (Santa Cruz, sc-13115), anti-alpha-tubulin (Sigma-Aldrich, T9026), anti-total-eNOS (BD Biosciences, #610296), anti-BNIP3 (Sigma-Aldrich, #B7931), anti-p62 (Sigma-Aldrich, P0067), anti-LC3 (Santa Cruz, #sc-2716291), anti-Parkin (Cell Signaling, #2132), Lamp1 (Invitrogen, #14-1071-82).

### Statistical analysis

Statistical analysis was performed using SPSS software ver. 18.0 (IBM, Armonk, NY, USA). The results are presented as mean ± SD. Depending on how many conditions were compared, either a two-tailed independent t-test analysis or one-way analysis of variance (ANOVA) with the Tukey’s post hoc analysis was conducted. The Pearson’s correlation coefficient (r) was used to measure the strength of a linear association between 2 variables. p<.05 was considered statistically significant for all analyses.

### Data availability

The data that supports the findings of this study are available from the corresponding author upon reasonable request.

### Author contributions

SGH and JYP conceived the study. SGH, JCS, JS, and SYC obtained the data. HW and JYP participated in data interpretation and discussed the results. SGH primarily drafted the manuscript and constructed figures. HW and JYP edited the manuscript. All authors edited the manuscript, and all authors approved the final submitted version.

## Acknowledgments

This work was supported by funding from the National Institute of Heart, Lung, and Blood Institute at the National Institutes of Health (R01 HL126952 to J.Y.P.); The American Heart Association Postdoctoral Fellowship grant (25POST1375863 to S.G.H) and the American Heart Association/Beatrice F. Nicoletti research grant (19POST34450157 to J.C.S).

## Competing Interests

The authors declare no competing interests.

---

**Supplemental figure 1.**
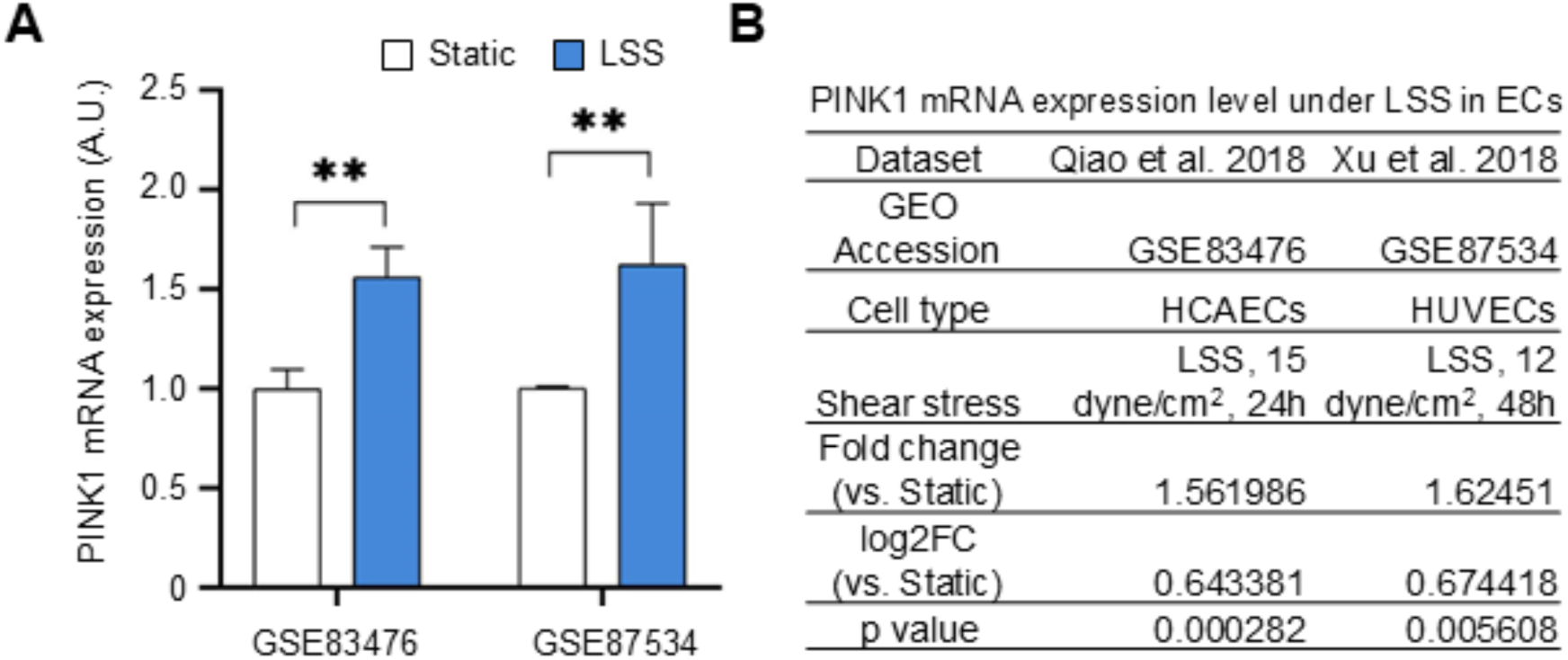
PINK1 mRNA expression in publicly-available RNAseq datasets. (A) PINK1 mRNA expression in RNAseq datasets from GSE83476 and GSE87534. (B) Detailed information of the RNAseq datasets. Data shown as means ± SD. **p<.01 by two-tailed independent Student’s t-test. A.U., Arbitrary Unit.

**Supplemental figure 2.**
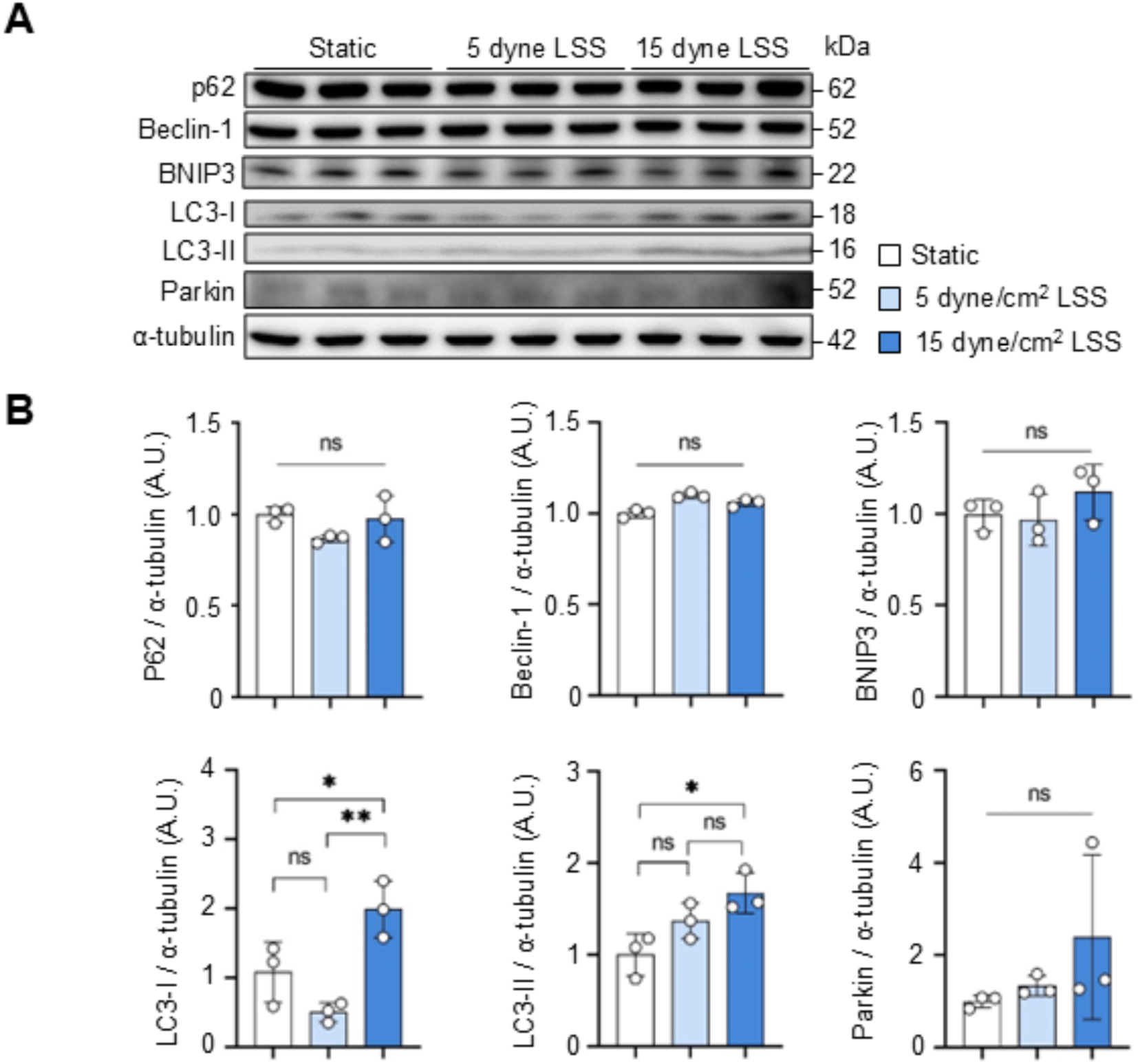
Mitophagy– and autophagy-related protein expressions under LSS. (A) Representative immunoblot images for the protein expression of p62, Beclin-1, BNIP3, LC3-I, LC3-II, and Parkin. α-tubulin was used as a loading control (n=3/per group). (B) Quantification plots for p62, Beclin-1, BNIP3, LC3-I, LC3-II, and Parkin protein expression. Data is shown as mean ± SD. ns, not significant. *p<.05. **p<.01 by one-way ANOVA followed by Tukey’s post hoc test.

## Notes

### Competing Interest Statement

The authors have declared no competing interest.

